# Isolation by environment is more important than isolation by distance along a tropical gradient in direct developing frogs

**DOI:** 10.1101/2023.06.01.543162

**Authors:** Ruth Percino-Daniel, Kara S. Jones, Thomas A. Maigret, David W. Weisrock, Daniel Piñero

**Affiliations:** Departamento de Ecología Evolutiva, Instituto de Ecología, Universidad Nacional Autónoma de México, Ciudad Universitaria, Coyoacán, Mexico City, Mexico CP 04510; Posgrado en Ciencias Biológicas, Universidad Nacional Autónoma de México, Coyoacán, Mexico City, 04510, Mexico; Department of Biology, University of Kentucky, Lexington, KY 40506, United States

**Author notes:** **Corresponding author**: Ruth Percino-Daniel. Departamento de Ecología Evolutiva, Instituto de Ecología, Universidad Nacional Autónoma de México.

**Keywords:** elevation gradient, *Craugastor*, environmental heterogeneity, isolation by environment, genomics, RADseq

## Abstract

Abiotic factors are important for defining population structure and limiting gene flow, especially in ectotherm species, the challenge is to identify which ones are the most important and can produce a pattern of isolation by environment. Our study aim is to quantify the extent to which divergence is driven by abiotic factors such as temperature, precipitation and elevation. To do so, we used a direct-developing frog species, *Craugastor loki* that occurs along a steep elevation gradient. Using restriction-site associated DNA sequencing (RADseq) from individuals collected from 100 m to 2250 m of elevation at 13 localities at Sierra Madre de Chiapas in southern Mexico we described population structure using a variety of model-based clustering and landscape genomics approaches. We found that populations sampled at higher elevation correspond probably to an undescribed new species of *Craugastor*, and that populations from *Craugastor loki* between 120 m and 1500 m are clustered in two different genetic groups: a Pacific slope group and a Central Depression slope group. We found signatures of isolation by environment more important that isolation by distance in contributing to genetic divergence in this group of frogs at a fine scale. The environmental variables such as: mean temperature of wettest and warmest quarter, annual mean temperature, and seasonality in temperature and precipitation play an important role on population differentiation. Our results underscore the importance of abiotic factors as drivers and highlight the use of different approaches to illuminate fine scale population divergence given the complexity of disentangling the contribution of both patterns of isolation.

## Introduction

Restriction of gene flow across a landscape can often be explained by the ubiquitous pattern of isolation by distance (IBD); (Wright, 1943), where genetic differentiation increases with the increase in geographic differences, and the differentiation can be accumulated by genetic drift. However, IBD assumes a uniform landscape and does not account for landscape heterogeneity. Gene flow can also be limited in environmentally heterogeneous landscapes due to divergent selection driving genetic differentiation, leading to isolation by environment (IBE). IBE can be identified by assessing the contribution of environmental factors on genetic variation while controlling for geographic distance (Wang, 2013; Wang & Bradburd, 2014). IBD and IBE can easily be confounded with one another and disentangling which of these processes is more important is key to understanding how the landscape influences genetic variation.

Recent studies in amphibians have shown that different climatic variables (e.g., annual mean temperature, isothermality, temperature range and precipitation) function as drivers of population divergence, thus producing a pattern of IBE (Medina et al., 2021; Páez-Vacas et al., 2022). Thermal differences can act as an environmental pressure on phenotypic traits through divergent selection and increase population differentiation (Páez-Vacas et al., 2022). Likewise, seasonal variation in temperature influences the movement and behavior of frogs (Seebacher et al., 2012) resulting in limited gene flow. Indeed, candidate single nucleotide polymorphisms (SNPs) and structural variants associated with temperature in frogs have been identified and suggest a potential functional role in life history strategies (Cayuela et al., 2021; Medina et al., 2021). Precipitation can limit amphibian distribution at both the regional and local scales (Ochoa-Ochoa et al., 2019). In dry seasons, frogs tend to move less because their ability to absorb water from the ground decreases when soil water potential decreases (Vitt & Caldwell, 2014; Wells, 2007). Despite an increasing number of studies that explore the contribution of environment on population divergence in ectotherm species, much remains to be explored in the way in which environmental variables shape gene flow and genetic diversity at a fine scale. This is particularly true in tropical forests, where the potential for biodiversity loss is higher.

At small scales, tropical mountains are highly heterogeneous in temperature, precipitation, and humidity (Stevens, 1992). This is especially clear in a landscape with an elevation gradient where climatic conditions vary with altitude. Thus, they are ideal for assessing the potential for IBE to limit gene flow (Funk et al., 2016) and to study which factors can lead adaptive differentiation and identify local adaptation (Luquet et al., 2015; Wake, 1987).

The Sierra Madre de Chiapas is a physiographic region with an ample variety of habitats, where lowland and intermediate habitats are influenced by the orientation of its slopes: the Pacific and the Central Depression. The Pacific slope is more humid in comparison to the Central Depression, due to a continental influence and a rain shadow effect. Furthermore, the region offers a natural elevation gradient on both slopes. Given the natural differences of temperature and precipitation, we can expect that IBE could contribute more than IBD to population structuring, especially for ectothermic organisms with low vagility and high site fidelity (Zamudio et al., 2016).

Here, we use the amphibian species *Craugastor loki* a direct-developing frog species found in North and Central America, inhabiting a variety of humid habitats in Chiapas, Mexico, as a model to investigate fine-scale genetic differentiation patterns. This group of frogs has a wide elevational distribution range extending from sea level up to 2200 meters above sea level (masl), particularly in southern Mexico (Frost, 2020; Streicher et al., 2014), and is often locally abundant. Consequently, this species is particularly suitable for investigating fine-scale genetic differentiation patterns across landscape-level gradients. Its continuous distribution allows for testing abiotic factors, such as the environment, as contributors to genetic differentiation and increased population structure. Our study aims to (1) quantify both fine-scale genetic differentiation and more divergent evolutionary relationships between populations of *C. loki* in the Sierra Madre de Chiapas, (2) test whether patterns of genetic variation in *C. loki* are best explained by geographic distance or environmental conditions, and (3) identify which specific environmental variables might be associated with patterns of genetic differentiation in *C. loki*.

## Materials and methods

### Study area and sampling

We collected 90 tissue samples of the direct-developing frog *C. loki* from 13 different localities in the Sierra Madre de Chiapas of southern Mexico (Figure 1, Table S1). Our sampling regime was designed to capture environmental variation along an elevation gradient, with sampling sites ranging from 120 to 2200 masl (Percino-Daniel et al., 2021). We collected mostly toe and, in some cases, liver tissues. Sampling was performed in the rainy season (June to November) in 2017. The number of samples varied from five to eight individuals per locality. We included *Craugastor pygmaeus* as an outgroup taxon, as it is closely related to the members of *Craugastor rhodopis* group, which includes *C. loki*. We also included *Craugastor rupinius*, which inhabits the Sierra Madre de Chiapas, is distantly related to *C. loki,* and is nested within the *Craugastor punctariolus* species group (Hedges et al., 2008).

**Figure 1.**
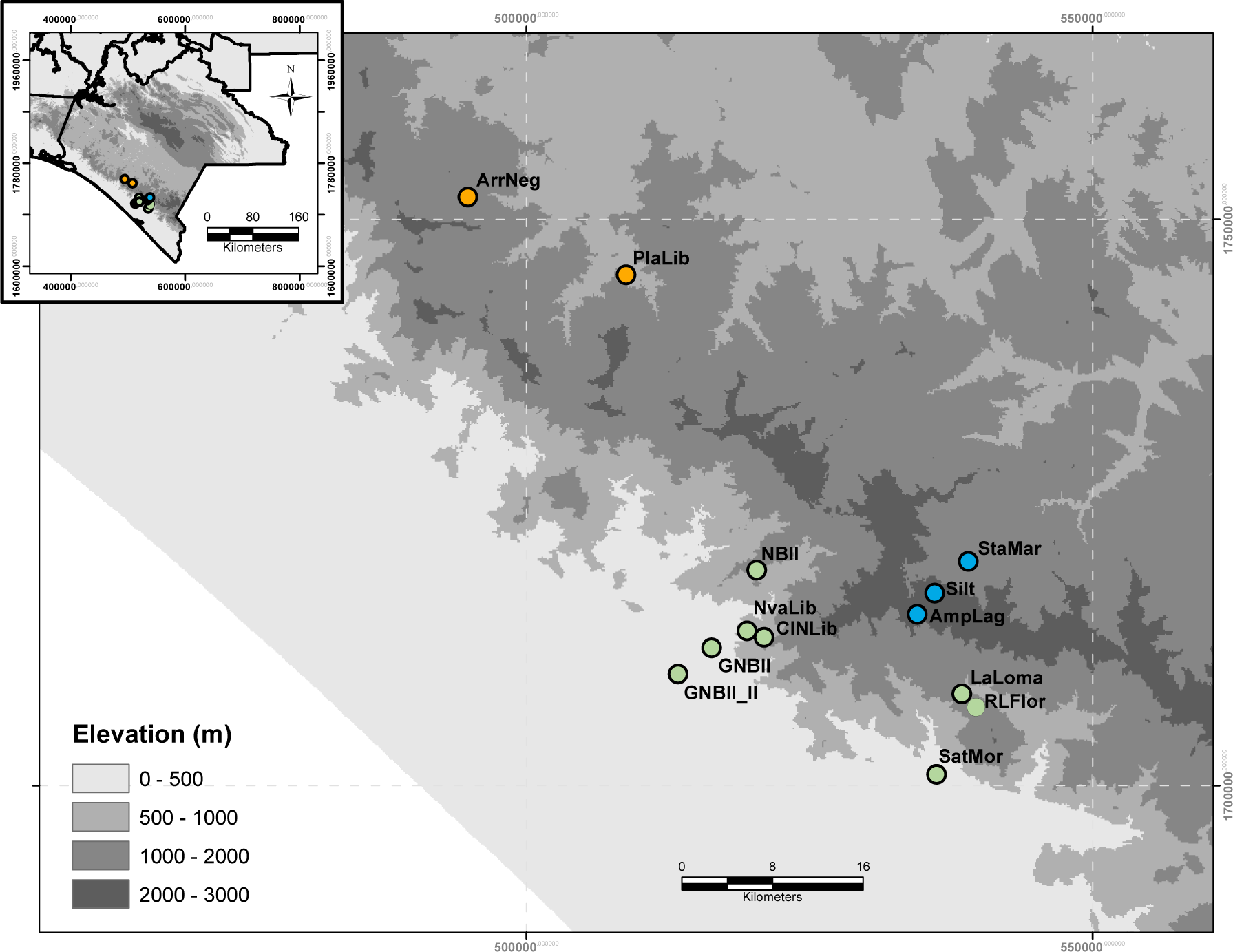
Sampled localities of *Craugastor* in Chiapas, Mexico. The orange and green circles correspond to the two genetic clusters of *C. loki* identified by Admixture and conStruct analysis, as seen in Figure 2. Blue circles correspond to the localities of *Craugastor* sp. from the highlands.

### DNA extraction and library preparation

Genomic DNA extractions were obtained using a QIAamp DNA Mini kit. Quality of extractions were checked with agarose gel electrophoresis and the quantity of extractions using a Qubit 3 Fluorometric Quantitation Kit (Invitrogen). DNA samples were standardized to (20ng/µl). We sequenced 90 *C. loki* samples using a restriction-site associated DNA sequencing (RAD-seq) method (Baird et al., 2008), employing the restriction enzyme *SbfI.* Library preparation and sequencing were performed at Floragenex Inc (Eugene, Oregon, USA). Sequencing libraries were run on two lanes of an Ilumina HiSeq 2000.

### Preprocessing and data assembly

We trimmed adaptors, filtered out low-quality data and demultiplexed raw data of the 90 samples using process_radtags in Stacks v.2.5. (Catchen et al., 2013). We subsequently assembled the data using Ipyrad v.0.7.30 (Eaton & Overcast, 2020), exploring parameter optimization with a focus on the maximum allowed divergence among allelic variants and the number of samples required per locus.

We made two datasets, the first (dataset 1) considered all samples from lowlands and intermediate elevation (120 to 1500 masl), the highlands (samples above to 2000 masl), plus the two outgroup samples of *C. pygmaeus* and *C. rupinius.* The second dataset (dataset 2) only included samples from lowlands to intermediate elevation (100 to 1500 masl).

Samples from higher elevations likely represent a cryptic species. For dataset 1, assembly and variant calling was performed using a clustering threshold of 0.85 and a minimum number of samples per locus of 35 (Supplementary Material Table S2). For dataset 2, we used 0.85 as clustering threshold and 30 as the minimum number of samples per locus. We filtered one SNP for each RAD-seq locus using VCFTOOLS v.4.2 (Danecek et al., 2011) and PLINK v.1.0.7 (Chang et al., 2015), and converted the SNP data to VCF format for landscape genomic analysis. Finally, we filtered dataset 2 allowing for at least 50% missing data, a minimum mean read depth of 26, and a minor allele frequency (MAF) of 0.01.

### Phylogenetic analyses

We reconstructed phylogenetic relationships among individuals of *C. loki* along the elevation gradient and outgroup species using dataset 1 and two tree-building methods. First, we used FastTree v.2.1.11 (Price et al., 2009) using nearest-neighbor interchange for the topology and subtree-prune and regraft moves (NNI + SPR) for branch length optimization. We used a general time-reversible nucleotide substitution model with a single rate per site (GTR + CAT). Second, we performed a maximum likelihood analysis in RAxML-HPC v.8.2.12 (Stamatakis, 2014) using a GTR + GAMMA nucleotide substitution model and saving the best-scoring tree. For node support we ran 100 bootstrap samples for both analyses. Trees were rooted with our choice of outgroup. Analyses were performed on the CIPRES Science Gateway v.3.3 (Miller et al., 2010)

### Non-spatial population structure analyses

We used two non-spatially aware methods to evaluate population structure. First, we used the non-parametric Discriminant Analysis of Principal Components (DAPC) with the Adegenet package v.2.1.1 in R v.3.6.1 (Jombart & Ahmed, 2011) to analyze both dataset 1 and dataset 2. To determine the number of the groups, we used the find.clusters function and selected the optimal *K* using the lowest BIC value exploring K 1 to 10. The optimal number of PCs to retain was selected using a cross-validation procedure (Jombart et al., 2010). Second, we used the model-based method Admixture v.13 (Alexander et al., 2009) to analyze dataset 2. We ran *K* from 1 to 5, with 10 iterations per *K* and selected the *K* with the lowest cross-validation error.

### Spatially informed population structure analyses

We used multiple approaches to investigate the relationship between geographic and genetic information. First, we inferred the strength of an IBD pattern and compared a spatial model with a non-spatial model using the R package conStruct v.1.3 (Bradburd et al., 2018) implemented in R. conStruct uses an explicit spatial component and models genetic variation in genotyped individuals as partitioned within or admixed specific number of discrete layers, within each layer the relatedness decays as a parameter function of the distance between samples (Bradburd et al., 2018). We analyzed *K* from 1 to 5, each with 10 replicates with 1000 MCMC iterations. We also used a cross-validation function to determine the statistical support for models with and without a spatial component. Because conStruct is sensitive to missing data, we collapsed some closely related individuals from the same population into a single, multi-individual samples (https://github.com/gbradburd/conStruct.git) and ran the cross-validation test using 35 multi-individual samples from 10 localities.

Next, we studied spatial and genomic variation across our sampled elevation gradient using a spatial Principal Component Analysis (sPCA; (Jombart, 2008) of dataset 2 implemented in Adegenet v.2.1.1. sPCA relies on an ordination approach for analyzing spatial and genetic patterns; using Moran’s *I* index, it identifies eigenvectors which maximize genetic variation and spatial autocorrelation and maps selected eigenvectors onto geographic space. Finally, we used the ResistanceGA v.4.0.14 software (Peterman, 2018) to associate specific landscape attributes with levels of gene flow. The landscape attributes we used were elevation, temperature, and precipitation (during the rainy season from July to November). ResistanceGA uses pairwise genetic dissimilarity and a genetic algorithm to optimize resistance surfaces. We employed least cost paths to estimate the effective distances across the landscape (Peterman, 2018) and used a linear mixed effects model with maximum likelihood to test for landscape effects, where pairwise genetic distance was the response variable and the optimized effective distance was the predictor variable. We used both approaches for each individual variable and for all combinations of the individual surface and null model (Peterman, 2018).

### Isolation by environment (IBE)

We tested for a signature of IBE using both temperature and precipitation variables, factors highly relevant to amphibian activity. Analyses were performed in the BEDASSLE v1.5 R package (Bradburd et al., 2013), which models the covariance of allele frequencies as a Gaussian process and uses a Bayesian model to estimate the contribution of environmental and geographic variables. For environmental variables, we obtained bioclimatic variables available to Mexico from Cuervo-Robayo et al. (2014) interpolated to ∼90m resolution.

Then, we ran a principal component analysis using the packages FactorMineR and factoextra in R v. 4.3.3 to summarize the 19 variables and to obtain the climatic variables that explained variance for the two principal components. The first dimension explained the 54.3%, which comprised three climatic variables: mean temperature of wettest quarter (Bio08), mean temperature of warmest quarter (BIO10), and annual mean temperature (Bio1), while the second dimension explained the 37.1 % was precipitation seasonality (Bio15) and temperature seasonality (Bio04) (Supplementary Material Figure S1, S2). Therefore, we used these five climatic variables as an environmental input matrix. Pairwise Euclidean distances were generated between sampling sites for environmental variables.

Pairwise great-circle geographic distances were generated between sampling sites using the function distm with the package geosphere in R. We standardized both distance matrices by dividing values by their standard deviation constants. We performed three replicate Markov chain Monte Carlo runs to ensure convergence of the parameters using the beta-binomial model, running one analysis for 5 million generations, and the second and third for 7 million generations, sampling every 1000 generations in all runs. Performance of the model was assessed by visualizing plot acceptance rates and parameter trace plots. We discarded the first 50% of samples as the burn-in and estimated the contribution of the environmental distance versus geographic distance to genetic differentiation using the ratio (*aE/aD*).

In addition, we used the multiple matrix regression with randomization analysis, with the package MMRR implemented in R (Wang, 2013) to quantify the contribution of the environmental and geographic distances on genetic differentiation. MMRR incorporates multiple regressions and can quantify how the genetic distances respond to changes in geographic and environmental distances, and provide a significance of each variable (Wang, 2013). MMRR provided significant values for regression coefficients (*β*) and coefficients of determination (*R^2^*), estimated by performing random permutations of rows and columns of the response variable, in this case, genetic differentiation and predictor matrices fixed, which in this case were the environmental matrix and geographic matrix independently (Wang, 2013). We performed 999 permutations in all calculations of significant values.

## Results

### Sequence data and bioinformatics

We obtained ∼661 million reads for the 94 samples, with 6.9 million mean reads per sample. After filtering, we obtained a total of 848 loci for dataset 1(Supplementary Material Table 2S) and 1801 loci for dataset 2 (Supplementary Material Table 3S). The assembled RADseq data set are available on Github repository as VCF files (https://github.com/rpercino/Landscape-genomics-Craugastor-).

### Phylogenetics and population structure

Both FastTree and RAxML phylogenetically placed samples from the highlands group (localities from elevation above 2000 masl, Figure 1) in a separate clade with 100% bootstrap support (blue shaded clade in Figure 2a), suggesting that they may belong to a different species of *Craugastor*. Some samples from intermediate elevations (samples from 1000 – 1200 masl) clustered with the outgroup samples: ArrNeg8 clustered with *C. pygmaeus* and the samples NBra1_1 and NBra1_4 grouped with *C. rupinius*. This is likely explained by the difficulty in differentiating adult members of *C. pygmaeus* from *C. loki*, and the difficulty in differentiating juveniles of *C. rupinius* from *C. loki*. Samples from the low-elevation localities and the majority of samples from intermediate-elevation localities were recovered as a clade of *Craugastor loki* in both analyses (orange and green shaded groups in Figure 2a).

**Figure 2.**
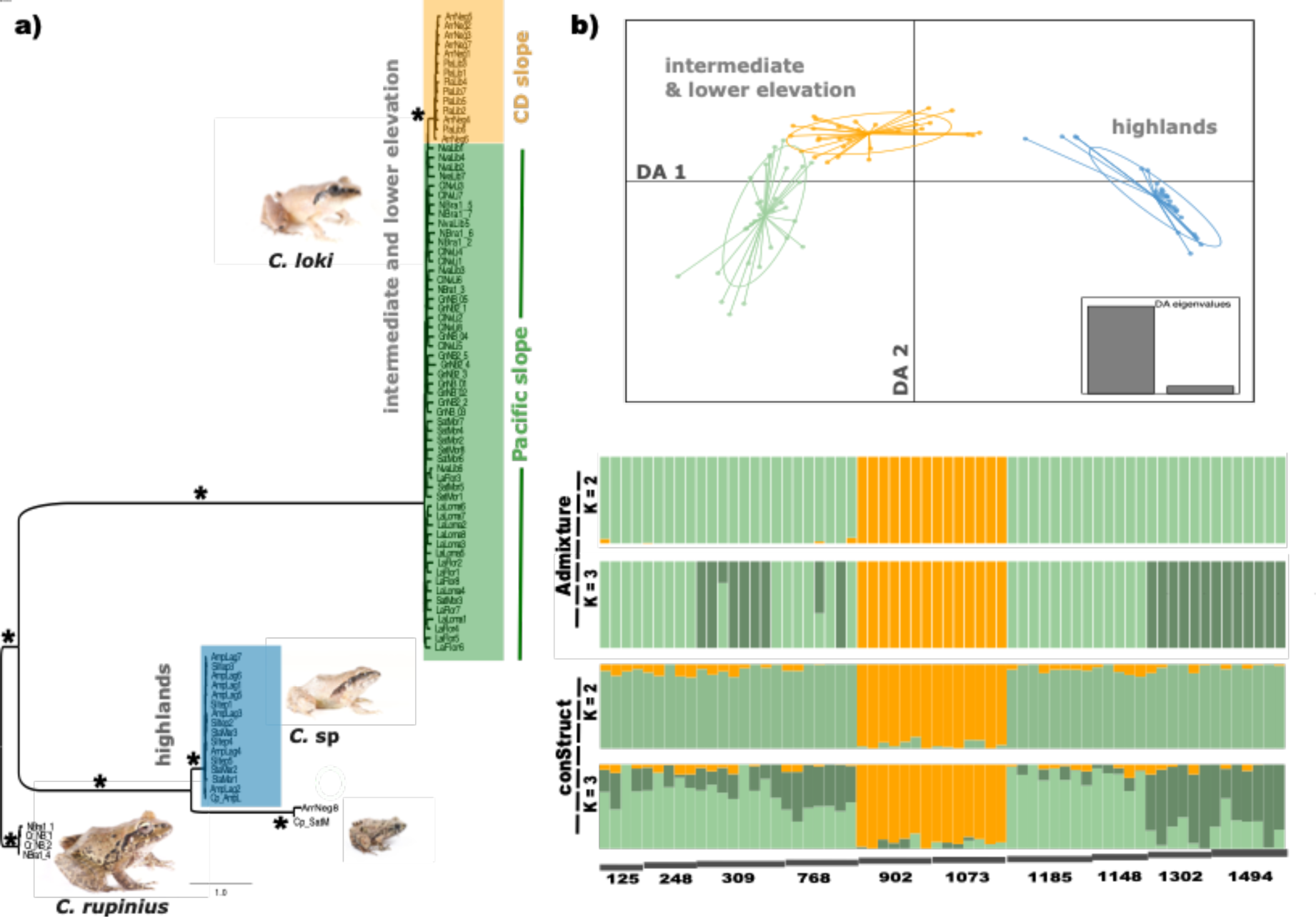
(a) Results from phylogenetic analysis using maximum likelihood performed in RAXML (asterisk indicates support bootstrap values >98). (b) discriminant analysis of principle components of the entire genomic data set. The individuals from higher elevation populations may correspond to other species of *Craugastor*. Colors correspond to the individuals sampled in the different localities shown in Figure 1. (c) Genetic assignment of the sampled populations using Admixture (from *K* = 2 to *K* = 3) and non-spatial model performed in conStruct. Individuals are ordered from lower to higher elevation. Elevation (in meters) is represented by horizontal bars below the assignment plots. Central Depression cluster and Pacific slope are shown in orange and green respectively. Photos in Figure 1 by R. Percino-Daniel (*C. rupinius* & C. sp) and J. E. Pérez Sanchez (*C. pygmaeus*).

The DAPC results for dataset 1 identified a *K* = 3 as the best-fitting model (Supplementary Material Figure S1a,b). The first axis accounted for 39.1% of the variation and identified the greatest separation between a cluster containing samples from the highlands, the outgroup samples (*C. pygmaeus* and *C. rupinius*), and the intermediate-elevation samples associated with those clades in the phylogenetic trees, and two clusters containing samples from the lowland and intermediate elevations (Figure 2b). Interestingly, four additional samples (GnNB2_4 LaFlor2, LaLoma1, LaLoma7) from intermediate elevations were placed closer to the “highland” cluster in ordination space. To verify the identity of these four samples, we reviewed fieldnotes and confirmed uncertainty in our taxonomic identification; thus, we omitted these individuals from dataset 2.

DAPC analysis of dataset 2 identified three groups, one of which is represented by the orange cluster that corresponds to the Central Depression slope (Supplementary Material Figure S1c). The other two groups (light green and dark green) identified in discriminant space are samples from the lowlands and intermediate elevations of the Pacific slope (Supplementary Material Figure S1c), accounting for 16.26% of the conserved variance. BIC scores stabilized at values of *K* = 3 (Supplementary Material Figure S1d).

Admixture analysis of dataset 2 identifying two groups with cross-validation support for *K* = 2 as the best fit model but additional structure was identified at *K* = 3 (Figure 2c, Supplementary Material Figure S4). One genetic cluster recovered samples from the Central Depression slope and the other clustered the lowland and intermediate samples from the Pacific slope (Figure 2c).

conStruct analyses recovered two genetic clusters (Figure 2c, Supplementary Material Figure S5): Central Depression slope (orange color) and Pacific slope (green color) including samples from low and intermediate elevations. However, there was no difference between non-spatial model and spatial model of conStruct (Supplementary Material Figure S6).

### Landscape genomics

The sPCA detected a significant pattern both globally (P = 0.003) and locally (P = 0.002). The first axis identified a break between the clusters (Supplementary Material Figure S7) associated with the Central Depression slope and the Pacific slope (Figure 3). However, some sampled localities fall at the boundary of the genetic break, corresponding to a lower elevation area.

**Figure 3.**
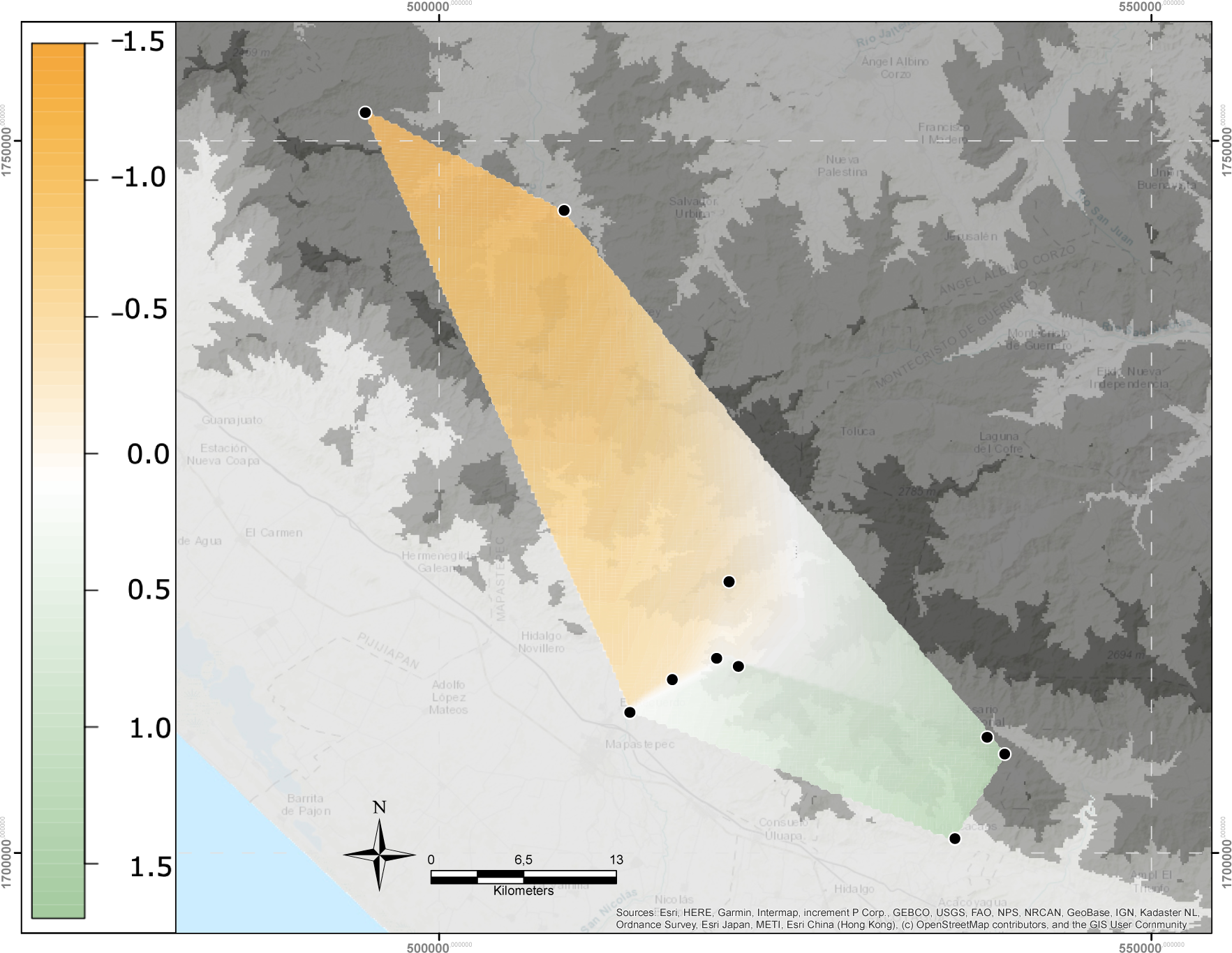
Interpolated spatial genetic structure based on sPCA superimposed over elevation map. The bar at left represents the interpolated vector scores, which corresponds to the degree of differentiation between individuals. The white color represents the break of the mountains edge of the highlands at Sierra Madre de Chiapas (lower elevation area). Black dots represent sampled sites.

The resistance surface analysis with ResistanceGA found the highest support for a model containing rainy season precipitation and elevation, although a competing model containing only temperature was nearly as well-supported (Table 1). Little to no support was found for a null model of no geographic structuring or for isolation-by-distance.

**Table 1.** Summary of model selection for the generalized linear mixed-effects models carried out in ResistanceGA. The null model assumes no geographic structure and the alternative models performed with environmental surfaces using temperature, precipitation, and elevation, where the interaction between elevation and precipitation is well fitted to the model.

| Layer | k | AIC | AICc | R2m | R2c | LL | delta.AICc | weight |
| --- | --- | --- | --- | --- | --- | --- | --- | --- |
| Elevation and<br>Precipitation | 7 | -7893.07 | -7891.07 | 0.0516 | 0.1276 | 3953.53 | 0 | 0.3965 |
| Temperature | 4 | -7891.19 | -7890.51 | 0.0425 | 0.1129 | 3949.60 | 0.5546 | 0.3005 |
| Precipitation<br>and<br>temperature | 7 | -7891.14 | -7889.14 | 0.0487 | 0.1247 | 3952.57 | 1.9287 | 0.1512 |
| Elevation and<br>temperature | 7 | -7890.51 | -7888.51 | 0.0486 | 0.1213 | 3952.25 | 2.5596 | 0.1103 |
| Precipitation | 4 | -7885.98 | -7885.30 | 0.0513 | 0.1389 | 3946.99 | 5.7669 | 0.0222 |
| Distance | 2 | -7884.71 | -7884.51 | 0.0260 | 0.0950 | 3944.35 | 6.5607 | 0.0150 |
| Elevation,<br>Precipitation,<br>and<br>temperature | 10 | -7885.42 | -7881.27 | 0.0494 | 0.1256 | 3952.71 | 9.7999 | 0.0030 |
| Elevation | 4 | -7880.71 | -7880.02 | 0.0260 | 0.0949 | 3944.35 | 11.0420 | 0.0016 |
| Null | 1 | -7836.83 | -7836.77 | 0 | 0.0630 | 3919.41 | 54.3006 | 6.41E-13 |

### Isolation by environment

Bayesian analysis of the relative contribution of IBE and IBD to genetic differentiation in BEDASSLE shows convergence in all runs producing similar results (Figure 4, Supplementary Material Figure S8), suggesting that IBE plays a role on genetic differentiation in *C. loki*. The mean temperature of wettest quarter (Bio08), mean temperature of warmest quarter (BIO10), and annual mean temperature (Bio1), precipitation seasonality (Bio15) and temperature seasonality (Bio04) show an important contribution on genetic differentiation than mere geographic distances. The *aE/aD* ratio indicates that the effective size of IBD would have equal the effect of IBE (Figure 4), that is, 370 Km of geographical distance would have the same effect of 1 degree Celsius of temperature variation or 1cm change in precipitation to have a similar effect on genetic divergence (Supplementary Material Table S4).

**Figure 4.**
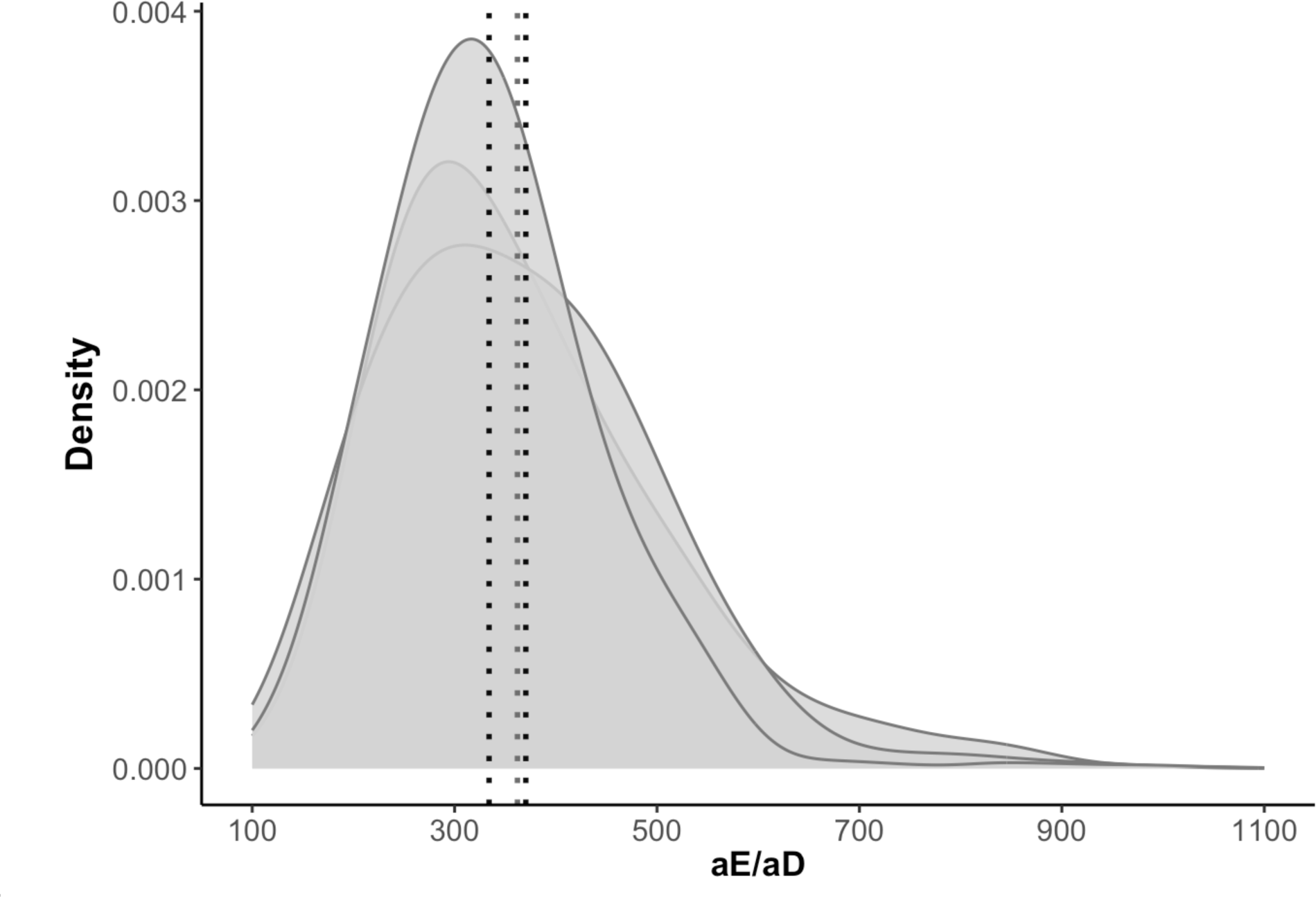
Results of the BEDASSLE analysis show the relative effect of the environmental distance (αE), using the following variables: mean temperature of wettest quarter (Bio08), mean temperature of warmest quarter (BIO10), and annual mean temperature (Bio1), precipitation seasonality (Bio15) and temperature seasonality (Bio04) and geographical distance (αD) on genetic differentiation. The mean ratio of *aE/aD* indicates the spatial distance (in Km) at which IBD would have equal the effect of IBE. For example, a value of 370 km of geographical distance would have the same effect of 1 degree Celsius variation of the temperature or 1cm change in precipitation. Each curve shows three independent runs.

The results using MMRR analysis show that both IBE and IBD play an important role in population genetic differentiation (Supplementary Material Figure S9). The regression coefficients for IBE were significant (*β* = 4.09, *P* = 0.014) as well IBD (*β* = 3.63, *P* = 0.005).

## Discussion

Disentangling which abiotic factors have an important role in population connectivity and limiting gene flow can be challenging, particularly given that abiotic environmental variables such as temperature and precipitation are usually autocorrelated with geography (Bradburd et al., 2013), and frequently also with elevation (Barry, 2008). Here, we found significant genetic differentiation between populations of the direct-developing frog *Craugastor loki,* and further found that patterns of gene flow were best explained by both patterns: isolation by environment as well by isolation by distance using a variety of approaches such as spatial and non-spatial analyses as Bayesian analyses and multiple matrix regression.

We found that *Craugastor loki* inhabits areas from sea level to ∼1500 masl. Previous work on the evolutionary relationship of the *Craugastor rhodopis* group (including *C. rhodopis, C. occidentalis* and *C. loki*) using mitochondrial DNA suggested three clades of *C. loki* occurring from Mexico to Central America, with the southern clade inhabiting mainly lowland areas (Streicher et al., 2014). Here we confirm that *C. loki* occupies mainly lowland and intermediate elevations and that samples above 1500 masl possibly correspond to another species. This group of frogs are highly polytypic and field identification is challenging due to a lack of diagnostic characters (Campbell & Savage, 2000; Crawford & Smith, 2005). Here, both phylogenetic and clustering results of our genomic data show that the members of *Craugastor* found above 1500 masl are relatively distinct from lower elevation populations of *C. loki*, indicating that they either belong to another species that we did not include in the analysis (e.g., *C. matudai*, *C. montanus, C. greggi*) or possibly are an unidentified species. Our sampling covers different localities that occur near the type locality of these species, like Cerro Ovando (Figure 1), the type locality where *C. matudai* (Taylor, 1941) and *C. montanus* occur (Taylor, 1942). Both species were described by museum specimens as *C. loki*. *Craugastor greggi* (Lynch, 1965) is also supposed to be present in this area, inhabiting cloud forest like the highland ecosystem where we sampled. Even if all those species represent different groups in the taxonomical classification of *Craugastor*, in the field these frogs could not to be distinguished from each other and the original descriptions did not describe the vivid coloration of live individuals. Further work and a geographic more extensive sampling are needed to clarify the evolutionary relationships among these taxa.

Within *Craugastor loki*, we identified two main groups: one cluster from the Central Depression slope and the other from the Pacific slope. Spatial analyses (e.g., sPCA) show a discontinuity in the area sampled that matches a valley that breaks the range of the Sierra Madre de Chiapas (Figure 3). It may be that the dispersal of frogs along the Sierra Madre de Chiapas could be restricted by this break in elevation. Frogs often have low dispersal capacity and limited movement up and down elevation gradients (Duellman & Trueb, 1994). In addition, it seems that the break identified by our spatial analysis is reflected in local forest communities. To the southeast of where the range breaks the local conditions become more humid. In contrast, to the northwest the conditions of the Sierra Madre de Chiapas are usually drier and more influenced by the Central Depression, resulting in a continuous dry forest habitat.

Furthermore, the ResistanceGA and IBE analysis (Bedassle and MMR) suggest an effect of climatic variables on spatial genetic differentiation. Temperature and precipitation are important factors influencing the natural history and ecology of these direct-developing frogs, characterized by the lack of a water-dwelling larval phase and their use of substrate humidity for reproduction (Duellman & Trueb, 1994). The Central Depression slope cluster presents a thermal landscape different from the Pacific slope, where the former is characterized mainly by temperature seasonality (Percino-Daniel et al., 2021). Hence, our results suggest that the abiotic variables considered in our study, temperature and precipitation, have an important role in the pattern of IBE.

Several studies on amphibians have documented patterns of genetic differentiation across elevation gradients (Funk et al., 2005, 2016) and have found evidence of phenotypic divergence associated with temperature and elevation, explained by a pattern of IBE (Medina et al., 2021). In addition, Medina et al. (2021) found a clinal pattern of genomic differentiation associated with temperature and identified some candidate SNPs associated with temperature and body size across the clinal gradient. Likewise, different studies of landscape genetics in amphibians also exhibit a common pattern of IBD given the limited dispersal capacities and fidelity to breeding sites (Chan & Brown, 2020; Nowakowski et al., 2017). Here, our results suggest a pattern of IBE and IBD are operating and shaping the population genetic differentiation at fine spatial scale. The climatic variables used here temperature and precipitation are important in general for amphibians, given that determine their body temperature and hence all their activities. In a previous study, Percino et al. (2021), studied some physiological traits from the same individuals of the species studied here. The frogs that inhabit at the Central Depression slope showed a different thermal sensitivity compared with the populations from Pacific slope. First, they exhibit high thermal accuracy, meaning that their body temperature is close to the microenvironmental temperature, and high thermal quality, that is, the thermal landscape offers suitable microhabitats with temperature close to body temperature. Thus, it is suggested that the landscape plays an important role on frog populations and eventually could drive phenotypic and probably genotypic divergence because of potential local adaptation.

Further work is necessary to specifically study the role of local adaptation or thermal plasticity coupled with evapotranspiration rates across the elevation of the Sierra Madre de Chiapas.

## Acknowledgments

We are very grateful to local authorities and landowners in Chiapas, who helped us and allowed us to work on their properties, and for their invaluable hospitality. Thanks to undergrad students of the UNICACH for their invaluable help in the fieldwork, especially to J. M. Contreras-Lopez. We are grateful to F. Mendez de la Cruz (IB-UNAM), for extending his collected permits to RP-D, issued by Secretaria del Medio Ambiente y Recursos Naturales (SEMARNAT), no. SGPA/DGVS/002490/18 (license no. FAUT-0074). We thank O. Godinez (CONABIO) for helping with environmental data and A. Cardenas for advice on some R scripts. We are grateful to the University of Kentucky Information Technology Department and the Center for Computational Sciences for granting access to the Lipscomb High Performance Computing Cluster resources, especially to Vikram Gazula. We also thank computer resources group from the CONABIO cluster, especially E. Campos for his advice. RP-D received a fellowship from Consejo Nacional de Ciencia y Tecnologia (CONACYT no. 434471) for her doctoral studies and grants for visiting scholar at University of Kentucky (CONACYT-Beca de Movilidad 2019) with additional support of the Programa de Apoyo a Estudios de Posgrado of UNAM (PAEP – UNAM 2019). This work was supported by the Instituto de Ecología, Universidad Nacional Autónoma de México to DP (IE-UNAM 2016-2019).

## Data availability statement

The data in vcf format are accessible to the repository in github at https://github.com/rpercino/Landscape-genomics-Craugastor-

## Author contributions

RP-D and DP designed the research. RP-D performed the fieldwork and lab work. RP-D, KSJ, TAM analyzed the data. DP, DW contributed with analytical tools. DP funding acquisition lead. RP-D lead the manuscript writing with input of all authors.

